# Alternative splicing of COQ-2 determines the choice between ubiquinone and rhodoquinone biosynthesis in helminths

**DOI:** 10.1101/2020.02.28.965087

**Authors:** June H. Tan, Margot Lautens, Laura Romanelli-Cedrez, Jianbin Wang, Michael R. Schertzberg, Samantha R. Reinl, Richard E. Davis, Jennifer N. Shepherd, Andrew G. Fraser, Gustavo Salinas

## Abstract

Parasitic helminths use two benzoquinones as electron carriers in the electron transport chain. In aerobic environments they use ubiquinone (UQ) but in anaerobic environments inside the host, they require rhodoquinone (RQ) and greatly increase RQ levels. The switch to RQ synthesis is driven by substrate selection by the polyprenyltransferase COQ-2 but the mechanisms underlying this substrate choice are unknown. We found that helminths make two *coq-2* isoforms, *coq-2a* and *coq-2e*, by alternative splicing. COQ-2a is homologous to COQ2 from other eukaryotes but the COQ-2e-specific exon is only found in species that make RQ and its inclusion changes the enzyme core. We show COQ-2e is required for RQ synthesis and for survival in cyanide in *C. elegans*. Crucially, we see a switch from COQ-2a to COQ-2e as parasites transition into anaerobic environments. We conclude that under anaerobic conditions helminths switch from UQ to RQ synthesis via alternative splicing of *coq-2.*

## Introduction

Parasitic helminths are major human pathogens. Soil transmitted helminths (STHs) such as the nematodes *Ascaris*, hookworms, and whipworms infect well over a billion humans and seven out of the 18 neglected diseases categorized by WHO are caused by helminths (CDC, 2019). Despite the huge impact on global health of these infections, there are few classes of available anthelmintics and resistance is increasing in humans and is widespread in some species that infect animals — for example, *Haemonchus contortus*, a wide-spread parasite in small ruminants, has developed multidrug resistance (Jackson, Coop, 2000, Kotze, Prichard, 2016). There is thus a serious need to develop new classes of anthelmintics that target the parasites while leaving their animal hosts unaffected. One potential target for anthelmintics is their unusual anaerobic metabolism which differs from that of their hosts. When STHs are in their free-living stages of their life cycles, they use the same aerobic respiration as their hosts and use ubiquinone (referred to here as UQ and as Q in other papers) as an electron carrier in their mitochondrial electron transport chain (ETC). However, when they infect their hosts, they encounter highly anaerobic environments. This is particularly the case when they live in the host gut and *Ascaris*, for example, can live for many months in this anaerobic environment (Dold, Holland, 2011). To survive, they use an alternate form of anaerobic metabolism that relies on the electron carrier rhodoquinone (RQ). Since hosts do not make or use RQ, RQ-dependent metabolism could be a key pharmacological target since it is required by the parasite but is absent from their mammalian hosts.

RQ is an electron carrier that functions in the mitochondrial ETC of STHs (Van Hellemond et al., 1995). RQ is a prenylated aminobenzoquinone that is similar to UQ (Figure 1), but the slight difference in structure gives RQ a lower standard redox potential than UQ (−63 mV and 110 mV, respectively) (Unden, Bongaerts, 1997, Erabi et al., 1976). This difference in redox potential means that RQ, but not UQ, can play a unique role in anaerobic metabolism. In aerobic metabolism, UQ can accept electrons from a diverse set of molecules via quinone-coupled dehydrogenases — these include succinate dehydrogenase and electron-transferring-flavoprotein (ETF) dehydrogenase. In RQ-dependent anaerobic metabolism, RQ does the reverse — it carries electrons to these same dehydrogenase enzymes and drives them in reverse to act as reductases that transfer electrons onto a diverse set of terminal acceptors (Tielens, Van Hellemond, 1998, van Hellemond et al., 2003). UQ thus allows electrons to enter the ETC via dehydrogenases; RQ can drive the reactions that let electrons leave the ETC and this difference in direction of electron flow is driven by the difference in redox potential (Figure 1A). This ability of RQ to provide electrons to an alternative set of terminal electron acceptors allows helminths to continue to use a form of mitochondrial ETC to generate ATP without oxygen. In this RQ-dependent anaerobic metabolism, electrons enter the ETC from NADH through complex I and onto RQ, and this is coupled to proton pumping to generate the proton motive force required for ATP synthesis by the F0F1ATPase (van Hellemond et al., 2003). The electrons are carried by RQ to the quinone-coupled dehydrogenases which are driven in reverse as reductases and the electrons thus exit onto a diverse set of terminal acceptors, allowing NAD^+^ to be regenerated and the redox balance to be maintained. This RQ-dependent metabolism does not occur in the hosts and it is thus an excellent target for anthelmintics (Kita, Nihei & Tomitsuka, 2003).

**Figure 1.**
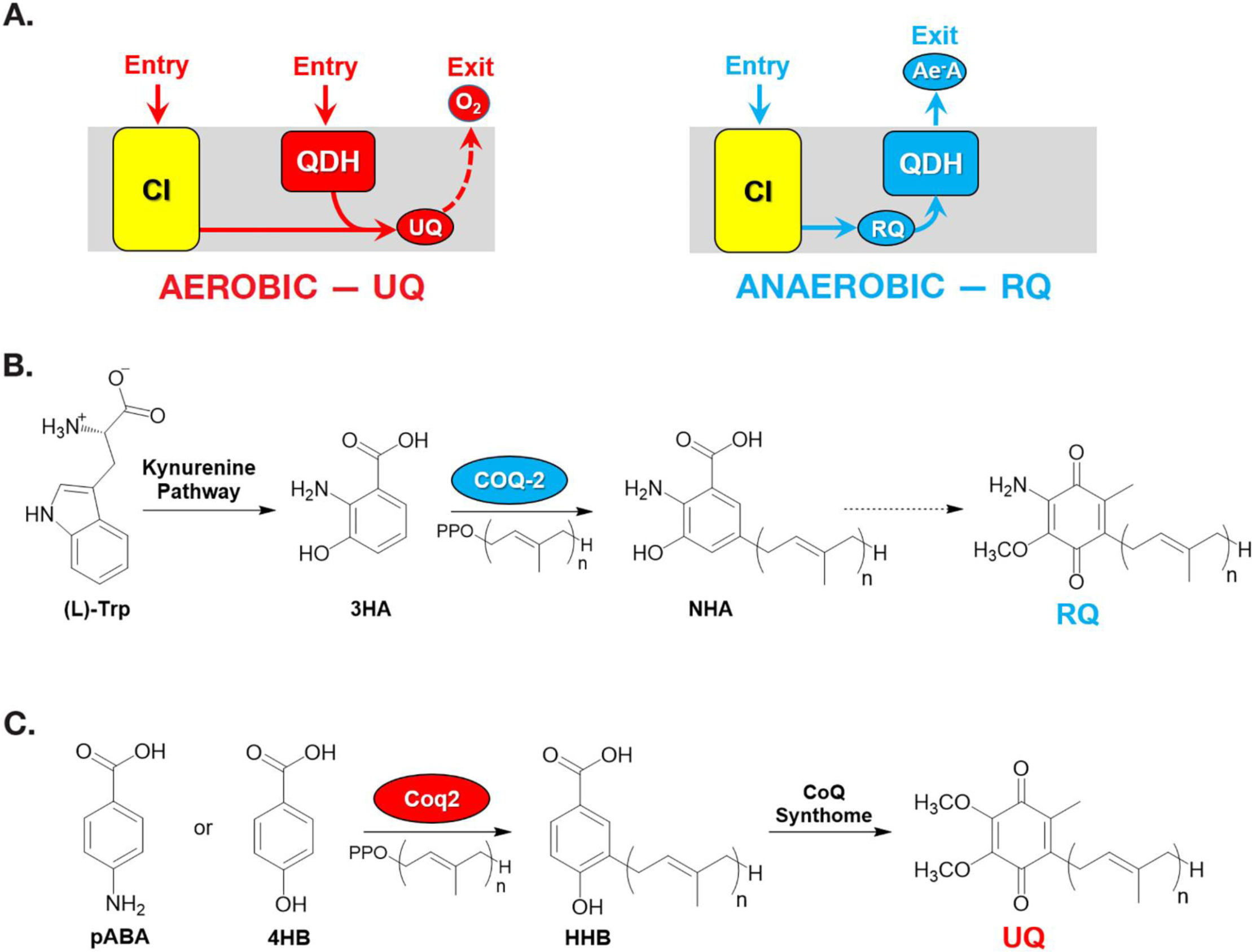
Rhodoquinone and ubiquinone biosynthesis and function in electron transport chains. (**A**) In aerobic metabolism, ubiquinone (UQ) shuttles electrons in the ETC from Complex I and quinone dehydrogenases (QDHs), such as Complex II and electron-transferring flavoprotein (ETF), which are ultimately transferred to oxygen. In anaerobic metabolism, rhodoquinone (RQ) reverses electron flow in QDHs and facilitates an early exit of electrons from the ETC at anaerobic electron acceptors (Ae^-^A), such as fumarate. (**B**) The RQ biosynthetic pathway in *C. elegans* requires *L*-tryptophan, a precursor in the kynurenine pathway. *L*-Trp is transformed into 3-hydroxyanthranilic acid (3HA) in four steps. It is proposed that 3HA is a substrate for COQ-2, producing 3-hydroxy-5-nonaprenylanthranilic acid (NHA) where n = 9. Transformation of NHA to RQ requires several shared proteins from the UQ biosynthetic pathway. (**C**) Yeast can use either *p*-aminobenzoic acid (pABA) or 4-hydroxybenzoic acid (4HB) as precursors to UQ. Prenylation is facilitated by Coq2 to form 3-hexaprenyl-4-hydroxybenzoic acid (HHB), where n = 6. Further functionalization occurs through a CoQ synthome (Coq3-Coq9 and Coq11) to yield UQ.

In the animal kingdom, RQ is present in several facultative anaerobic lineages that face environmental anoxia or hypoxia as part of their lifecycle. Among animals, RQ has only been described in nematodes, platyhelminths, mollusks and annelids (Van Hellemond et al., 1995). Key steps of RQ biosynthesis in animals were recently elucidated (Roberts Buceta et al., 2019, Del Borrello et al., 2019). In contrast to bacteria and protists, where RQ derives from UQ (Brajcich et al., 2010, Stairs et al., 2018), RQ biosynthesis in animals requires precursors derived from tryptophan via the kynurenine pathway (Figure 1B). Studies performed by two teams in *C. elegans* have recently demonstrated that animals that lack a functional kynureninase pathway (e.g. strains carrying mutations in *kynu-1*, the sole kynureninase) are unable to synthesize RQ (Roberts Buceta et al., 2019, Del Borrello et al., 2019). It is presumed that 3-hydroxyanthranilic acid (3HA here, also sometimes referred to as 3HAA) from the kynurenine pathway is prenylated in a reaction catalyzed by COQ-2. The prenylated benzoquinone ring can then be modified by methylases and hydroxylases (COQ-3, COQ-5 and COQ-6) to form RQ. This proposed pathway is analogous to the biosynthesis of UQ from 4-hydroxybenzoic acid (4HB) or *para*-aminobenzoic acid (pABA) (Figure 1C). The key insight from these previous studies is that the critical choice between UQ and RQ synthesis is the choice of substrate by COQ-2 — if 4HB is prenylated by COQ-2, UQ will ultimately be made, but if 3HA is used, the final product will be RQ. In most parasitic helminths there is a major shift in quinone composition as they move from aerobic to anaerobic environments e.g. RQ is less than 10% of *H. contortus* total quinone when the parasite is in an aerobic environment but is over 80% of total quinone in the anaerobic environment of the sheep gut and similar shifts occur in other parasites (Van Hellemond et al., 1995, Sakai et al., 2012, Luemmen et al., 2014). Somehow, COQ-2 must therefore switch from using 4HB to using 3HA as a substrate but the mechanism for this substrate switch is completely unknown. Understanding this mechanism is important — if we could interfere pharmacologically with the switch to RQ synthesis, it could lead to a new class of anthelmintics.

In this study, we reveal that two variants of COQ-2, derived from alternative splicing of mutually exclusive exons, are the key for the discrimination in the RQ/UQ biosynthesis. The removal of one of the mutually exclusive exons, present only in species that synthesize RQ, abolishes RQ biosynthesis in *C. elegans*. The analysis of COQ-2 RNA-seq data from parasites revealed that the RQ-specific exon expression is increased in hypoxic lifestages, while the alternative exon is increased in normoxic lifestages. We thus conclude that the alternative splicing of COQ-2 is the key mechanism that regulates the switch from UQ to RQ synthesis in the parasite life-cycle. We also propose that the reason that RQ is only synthesised in helminths, annelids, and molluscs is due to the independent evolution of COQ-2 alternative splicing in these animal lineages.

## Results

### The *C. elegans* COQ-2 polyprenyl transferase required for quinone biosynthesis has two major alternative splice forms

Our groups previously showed that if COQ-2 uses 4HB as a substrate, this will lead to synthesis of UQ; however, if it uses 3HA this will ultimately yield RQ. As parasites move from aerobic environments to the anaerobic niches in their host, they change their quinone composition from high UQ to high RQ. For this to occur, COQ-2 must switch its substrate from 4HB to 3HA but the mechanism for this switch is unknown.

We identified two distinct splice forms of *C. elegans coq-2*: *coq-2a* and *coq-2e*. These are annotated in the genome and confirmed by RNA-seq, by nanopore sequencing, and by targeted validation studies (Ramani et al., 2011, Kuroyanagi, Takei & Suzuki, 2014, Roach et al., 2020, Li et al., 2020). These two isoforms differ by the mutually exclusive splicing of two internal exons (6a and 6e), both of 134 nucleotides (see Figure 2A). We note that mutually exclusive splicing of cassette exons is very rare in the *C. elegans* genome and fewer than 100 such splicing events have ever been identified (Ramani et al., 2011, Kuroyanagi, Takei & Suzuki, 2014). Both *coq-2a* and *coq-2e* splice forms are abundant in *C. elegans* across all stages of development, where ∼30-50% of *coq-2* is the *coq-2e* isoform (Ramani et al., 2011, Gerstein et al., 2010, Grun et al., 2014).

**Figure 2.**
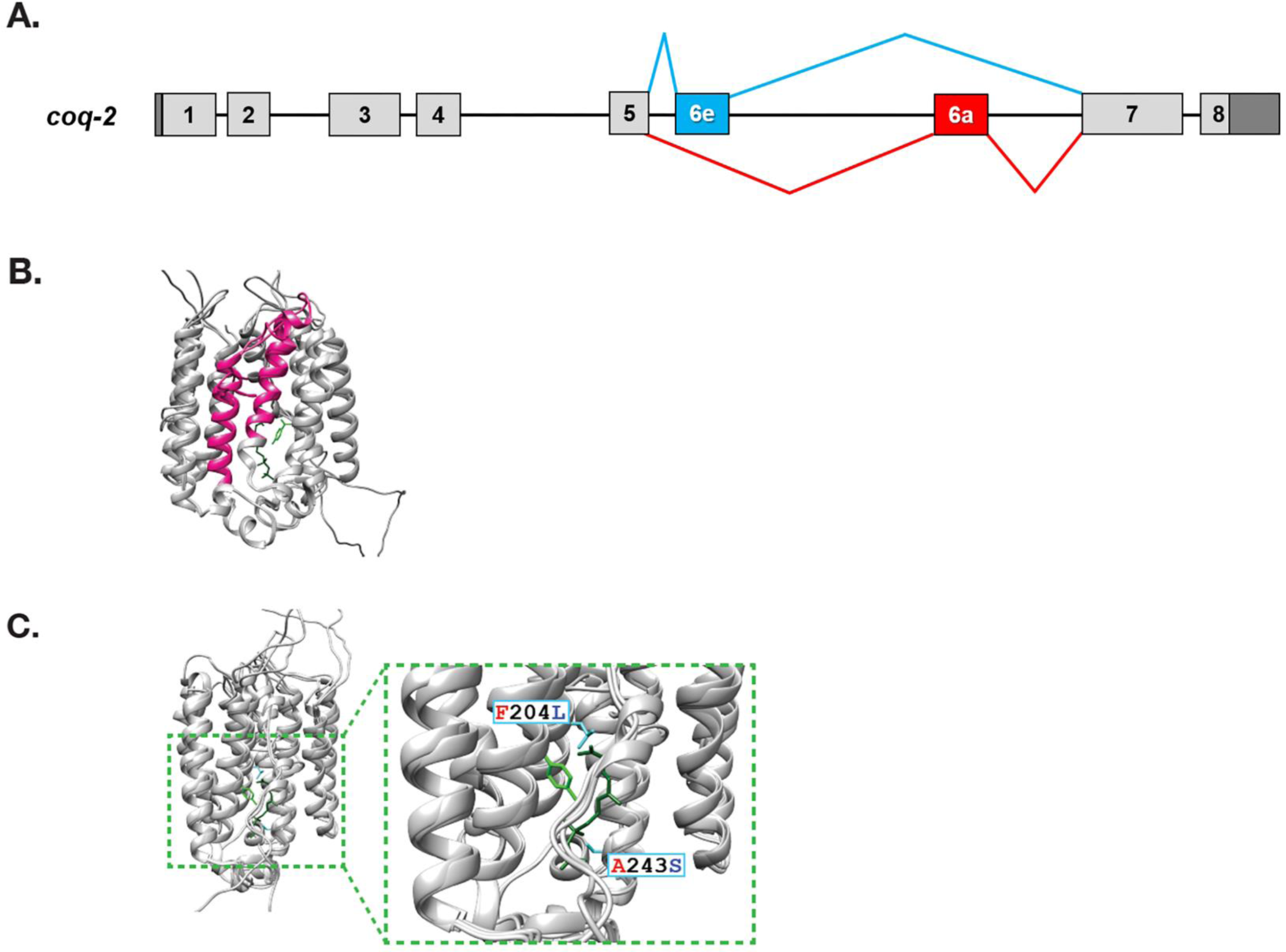
*C. elegans coq-2* gene model. (**A**) The *coq-2* gene contains two mutually exclusive exons, 6e (blue box) and 6a (red box), that are alternatively spliced (blue and red lines, respectively) generating two COQ-2 isoforms. Light grey boxes represent coding sequences of exons 1-5 and 7-8, black lines represent introns, and dark grey boxes denote 5’and 3’ untranslated regions of exons 1 and 8. (**B**) Alternative splicing of COQ-2 changes the enzyme core. The sequences of *C. elegans coq-2a* and *coq-2e* were threaded onto the crystal structure of the apo-form of the *A. pernix* COQ-2 homologue (PDB: 4OD5) in Chimera using Modeller. The region switched by mutually exclusive alternative splicing is shown magenta. (**C**) The alternative exons found in all RQ-synthesising species have two residues that are invariant (L204 and S243 show in cyan; *C. elegans* numbering) that are near the binding site of the two substrates. Substrates are the polyprenyl tail (dark green; geranyl S-thiolodiphosphate in the crystal structure), and the aromatic ring (light green; *p*-hydroxybenzoic acid (4HB) in the crystal structure). Note that COQ-2 is rotated from panel B to panel C for clarity.

To examine how the alternative splicing of COQ-2 might affect its function, we threaded the predicted COQ-2a and COQ-2e protein sequences onto the solved crystal structure of the archaean *Aeropyrum pernix* COQ-2 orthologue (PDB:4OD5). We found that the splicing change causes a switch in two alpha-helices at the core of the COQ-2 structure (Figure 2B). This is a region of the protein that is thought to form a hydrophobic tunnel along which the aromatic substrates must pass to the active site for the key polyprenylation reaction (Desbats et al., 2016) suggesting that the change in splicing could affect COQ-2 substrate selection and thus could explain a shift from UQ to RQ synthesis. We therefore examined whether similar COQ-2 alternative splicing is seen in parasitic helminths and how the different splice forms compare to COQ-2 sequences in parasite hosts which do not make RQ.

### Parasitic helminths have a distinct splice form of *coq-2* that is not present in any of the parasitic hosts

*C. elegans* has two major isoforms of *coq-2* which are the result of mutually exclusive alternative splicing of two internal exons and affects the core of the enzyme. If this alternative splicing affects the choice of COQ-2 substrate and thus the switch from UQ to RQ synthesis, we reasoned that parasitic helminths that make RQ should have a similar gene structure and that the hosts that do not make RQ should not. We used both gene predictions and available RNA-seq data to examine the *coq-2* gene structure and splicing in parasitic helminths and their hosts (human, sheep, cow, and rat respectively). As shown in Figure 3, all the parasites examined have a similar *coq-2* gene structure with the same two mutually exclusive internal exons — note that while many of the parasite gene structures were not correctly annotated, all the relevant exon junctions in Figure 3 were manually annotated and confirmed with RNA-seq data (see Materials and Methods). Remarkably, we find a similar gene structure with the same mutually exclusive exons in all parasites known to make RQ. Furthermore, annelids and molluscs are the only other phyla known to make RQ and their *coq-2* orthologues also show mutually exclusive alternative splicing of homologous exons. This gene structure is only seen in species that make RQ — we find no evidence in any available data for similar alternative splicing in any mammalian hosts (human and mouse are shown as representatives in Figure 3) or in other lineages that lack RQ, such as yeasts. This suggests that this *coq-2* alternative splicing could indeed be linked to the ability to synthesize RQ.

**Figure 3.**
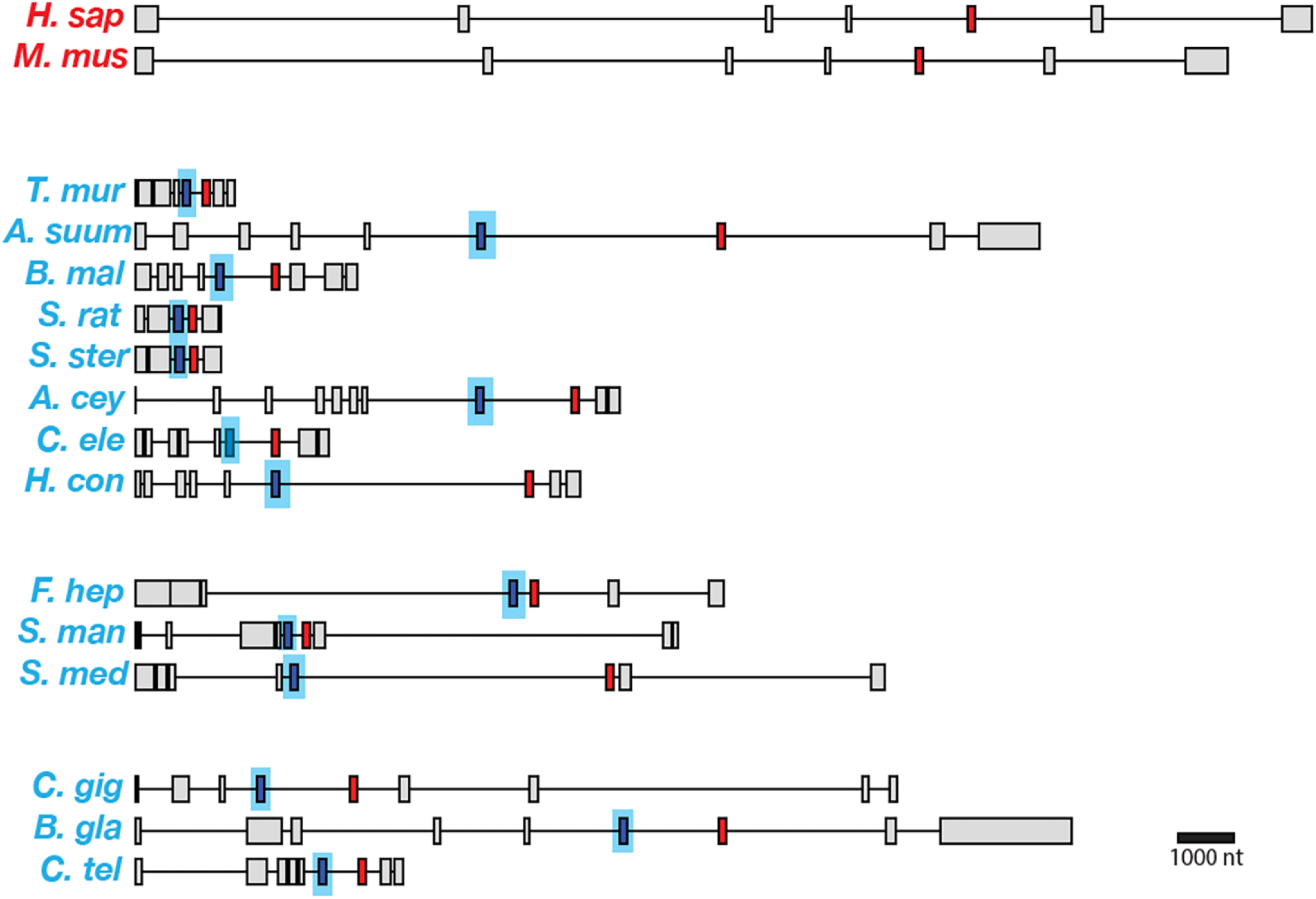
Gene models for *coq-2* orthologues in various species. Parasitic helminths, as well as annelids and molluscs, have 2 internal exons that are spliced in a mutually exclusive manner. In contrast, humans and other hosts only express one exon that is homologous to exon 6a of *C. elegans coq-2*. The a-form exon (red) shares greater similarity to the exon present in species that do not make RQ, while the e-form (blue) is present only in RQ-producing species. The genes used for each species are listed in Supplemental Table 3. The gene structures shown are based on genome annotations but in many cases include manual reannotations — in all such cases, the manual annotations are confirmed with RNA-seq data.

We aligned the two mutually exclusive exons across helminth species and compared them to the similar regions of their host COQ-2 sequences, and of other eukaryotes that cannot make RQ (*S. cerevisiae, S. pombe*) (Figure 4). We find that the *coq-2a*-specific exon is similar to the pan-eukaryotic COQ-2 sequence whereas the *coq-2e*-specific exon is distinct in all species that make RQ. We examined the alignments of the a-specific and e-specific exons and identified two residues that are strictly conserved in pan-eukaryotic COQ-2 sequences, Phe204 and Ala243 (*C. elegans* numbering), that are switched to a Leu and a Ser residue in all COQ2-e-specific exons that we examined (Figure 4). These residues sit very close to the substrates in the active site of the enzyme (Figure 2C) and we note that mutation of the equivalent Ala243 residue dramatically affects the ability of human COQ2 to make UQ (Desbats et al., 2016). Altogether these results suggest that animals that make both UQ and RQ make two forms of COQ-2 — one looks similar to that in all other eukaryotic species, whereas the other has a single exon that appears to be specific for species that make RQ. To test whether these two COQ-2 isoforms have distinct roles in UQ and RQ synthesis, we turned to *C. elegans.*

**Figure 4.**
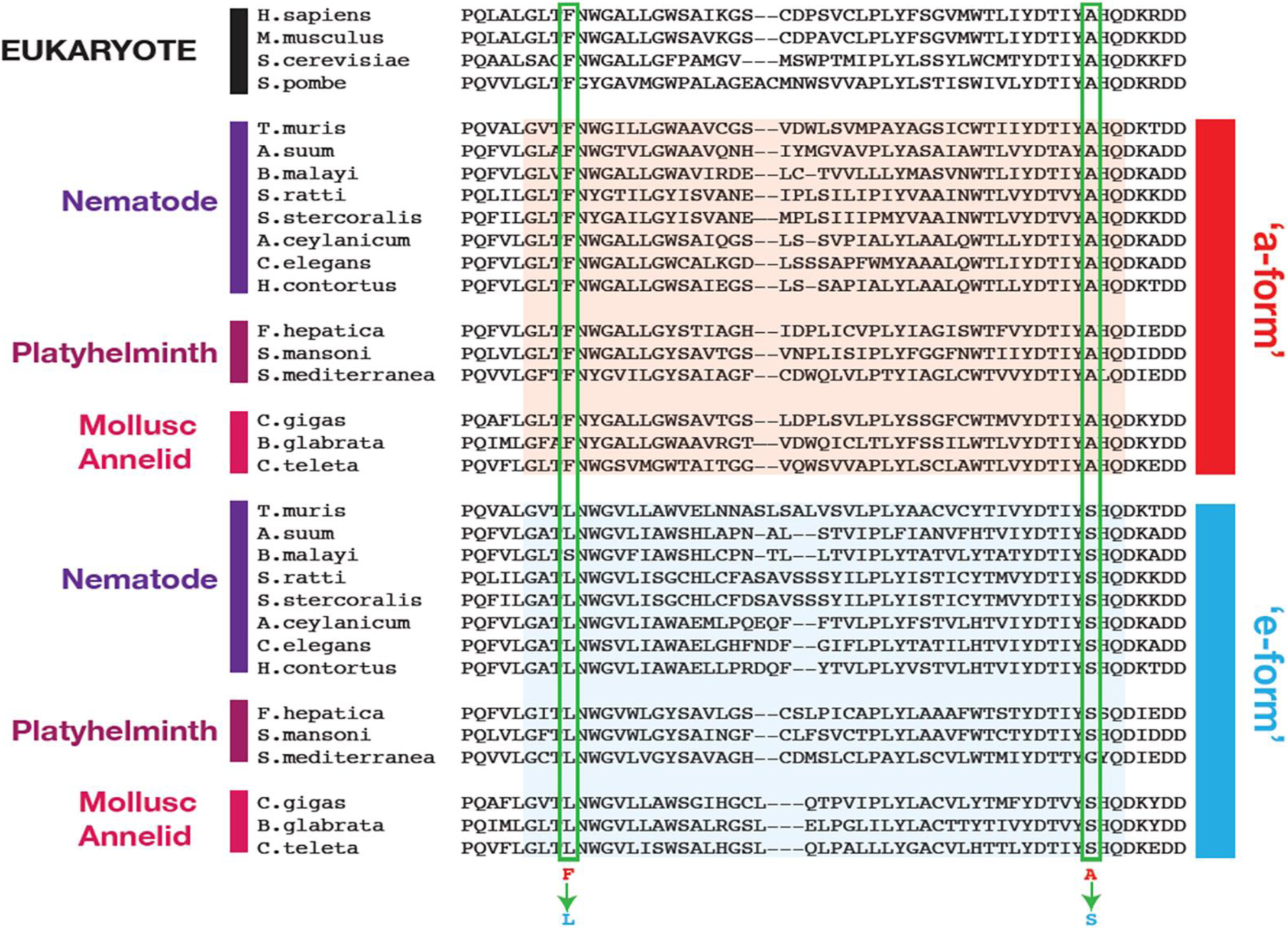
Conserved changes between a- and e-form exons across RQ-producing phyla. Amino acid sequences of COQ-2 orthologues were aligned using Clustal Omega (Madeira et al., 2019). The sequences of exons homologous to exon 6a/e in *C. elegans*, as well as the flanking 5 amino acid sequences were used to generate the alignment. Sequences of the mutually exclusive exons are shaded in red (a-form) or blue (e-form). Two residue changes between the a- and e-forms are highlighted (Phe to Leu, Ala to Ser) and are invariant across diverse species that make RQ. The COQ-2 orthologues and exons used for each species are listed in Supplemental Table 3.

### RQ synthesis requires the *coq-2e* isoform

We found that helminths make two major isoforms of *coq-2* whereas their hosts only make a single isoform. To test the requirement for each of the two major helminth isoforms of *coq-2* for RQ synthesis, we used CRISPR engineering to generate *C. elegans* mutant strains that either lack *coq-2* exon 6a (*coq-2(syb1715)*) or *coq-*2 exon 6e (*coq-2(syb1721)*) — we refer to these as *coq-2*Δ6a and *coq-2*Δ6e respectively from here on (see Figure 5A and Supplemental Table 1 for details of engineering). We find that the *coq-2*Δ6e strain makes essentially no detectable RQ but has higher levels of UQ, whereas *coq-2*Δ6a has greatly reduced UQ levels but higher RQ levels (Figure 5B, Supplemental Table 2). We conclude that the *coq-2e* isoform, that includes the helminth-specific exon 6e, is required for RQ synthesis.

**Figure 5.**
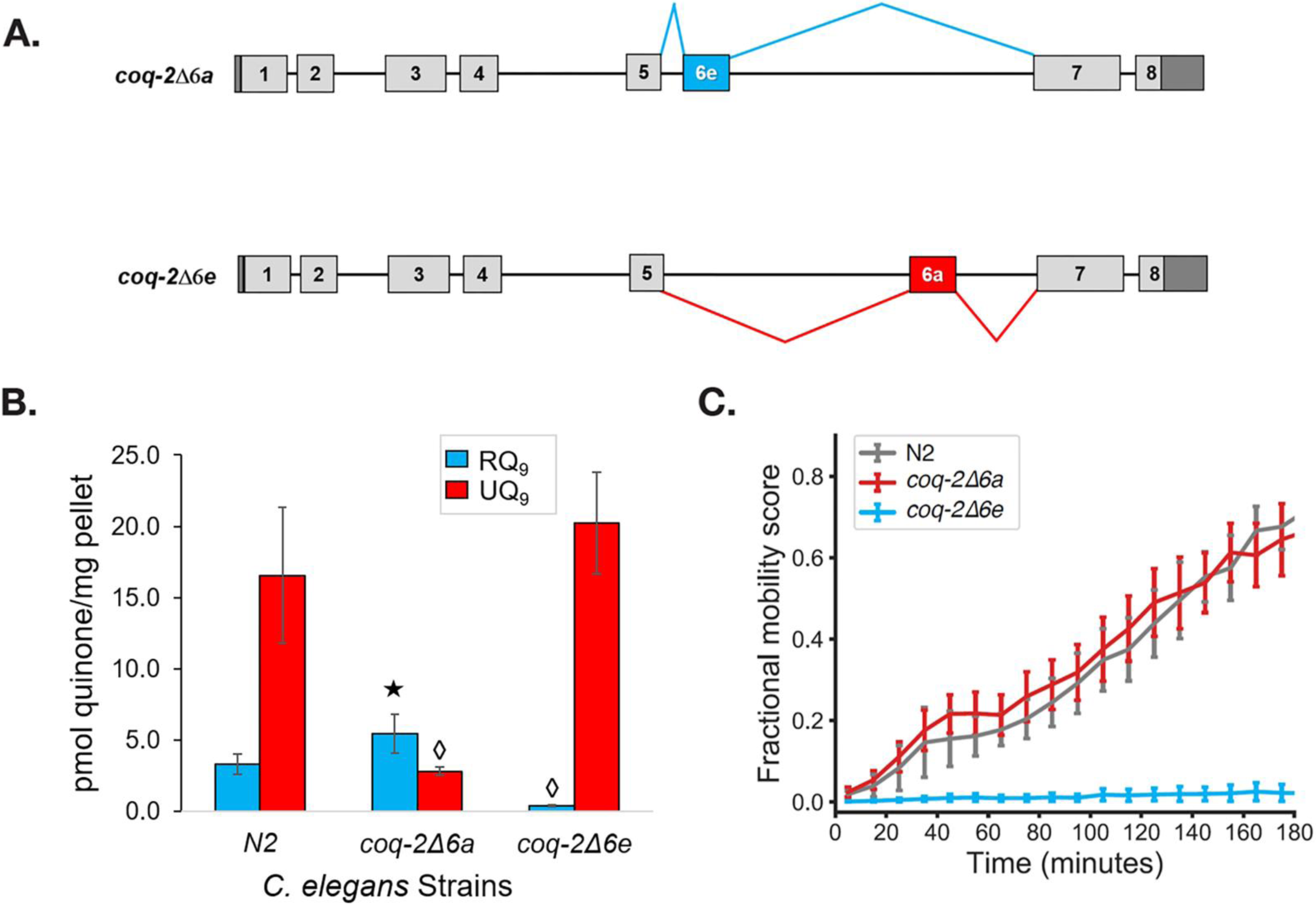
The *C. elegans coq-2* edited strains and effects of exon 6a and 6e deletions on quinone biosynthesis. (**A**) Mutant strains were generated in *C. elegans* by deletion of exon 6a (*coq-2*Δ6a) or exon 6e (*coq-2*Δ6e). (**B**) Deletion of exon 6a from the *coq-2* gene significantly increased the level of RQ_9_ (p = 0.013) and significantly decreased UQ_9_ (p < 0.001) compared to the N2 control. In contrast, the deletion of exon 6e decreased RQ_9_ to a negligible level (p < 0.001) and slightly increased the level of UQ_9_ (p = 0.130) compared to N2. Statistically significant increases and decreases with respect to N2 levels are denoted with ⋆ and ◊, respectively; error bars reflect standard deviation where N =4. (**C**) Deletion of *coq-2* exon 6e affects the ability of worms to survive extended KCN treatment. Wild-type (N2) and *coq-2* mutant L1 worms were exposed to 200 µM KCN for 15 h. KCN was then diluted 6-fold and worm movement was measured over 3 h to track recovery from KCN exposure (see Materials and Methods). Worms without exon 6e could not survive extended treatment with KCN while deletion of exon 6a had little effect on KCN survival. Curves show the mean of 4 biological replicates and error bars are standard error.

To further examine whether COQ-2e is required for RQ synthesis and thus for RQ-dependent metabolism, we tested whether the *coq-2*Δ6a and *coq-2*Δ6e strains could survive long-term exposure to potassium cyanide (KCN) (Figure 5C). We previously showed that when *C. elegans* is exposed to KCN it switches to RQ-dependent metabolism and that while wild-type worms can survive a 15 h exposure to KCN, *C. elegans* strains that do not make RQ cannot survive. We found that while the *coq-2*Δ6a strain (that can make RQ) survives 15 h of KCN exposure as well as wild-type animals, the *coq-2*Δ6e strain that makes no RQ does not survive, confirming the functional relevance of the *coq-2e* isoform as being critical for RQ synthesis.

### Regulation of the alternative splicing of *coq-2* in helminths

Helminths make two isoforms of COQ-2 — COQ-2a resembles the pan-eukaryotic consensus and cannot make RQ, whereas COQ-2e includes an exon that is only found in species that make RQ and COQ-2e is required for RQ synthesis. Changing the levels of *coq-2a* and *coq-2e* splice forms could thus regulate the switch from UQ synthesis in the aerobic environment outside the host to RQ synthesis in the host gut. We thus examined RNA-seq data to see whether parasites switch from between these isoforms as they switch between these environments.

*Ascaris suum* has a relatively simple life cycle (schematic Fig 6A) and is the pig equivalent of *Ascaris lumbricoides* which infects ∼900 M humans. Eggs are laid in the host and emerge via defecation and the L1-L3 larval stages develop within the egg outside the host. The L3 infective larval stage then enters the host via ingestion into the digestive tract. These then leave the digestive tract and make their way to the lungs where they develop into L4 larvae and, finally, the L4 re-enter the digest tract and move to the small intestine where adults develop.. The adults remain in the anaerobic environment for the remainder of their life. The free-living larval stages have relatively low RQ (∼ 35% of total quinones), whereas in adults, RQ is ∼ 100% of total quinones (Takamiya et al., 1993). We used RNA-seq data (Wang et al., 2011, Wang et al., 2012) to analyse *coq-2* isoforms in free-living stages and in the adults to examine whether there was switch from *coq-2a* to *coq-2e* as the parasites switch from low RQ aerobic-respiring embryos and larvae to high RQ anaerobic adults. We see a clear switch: ∼60% of *coq-2* transcripts are *coq-2e* in the free-living, aerobic stages but >90% is the RQ-synthesising *coq-2e* form in adults (Figure 6B). Increased *coq-2e* levels thus correlate with increased RQ levels.

**Figure 6.**
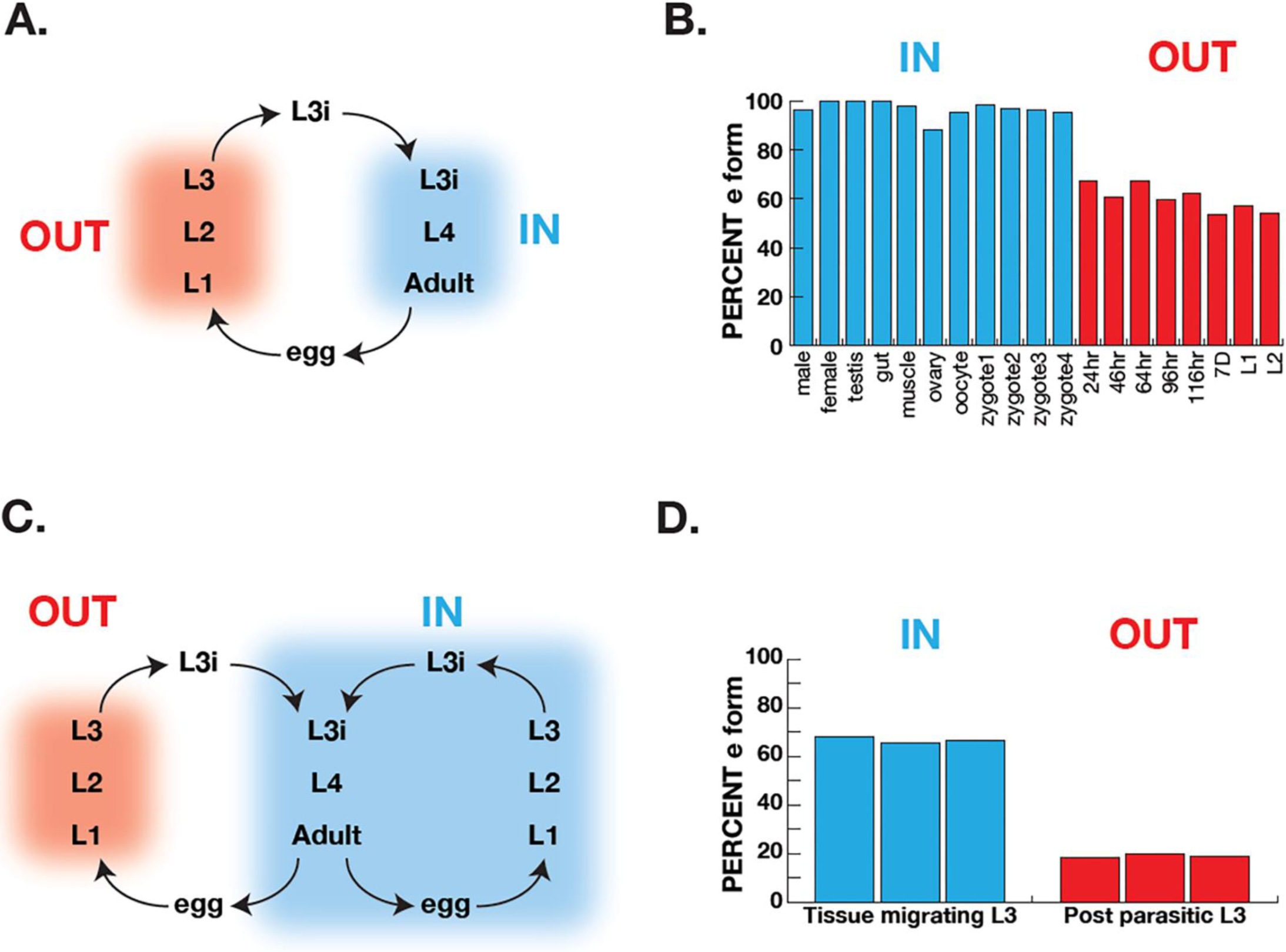
Correlation of COQ-2 splicing with change from aerobic to anaerobic life stages. (**A**) Schematic of life cycle of *A. suum*. ‘OUT’ denotes aerobically-respiring free-living stages; ‘IN’ indicates stages living inside the host intestine. (**B**) Graph indicates the percentage of all COQ-2 transcripts that include the RQ-specific exon (‘Percent e-form’) in a number of life cycle stages, sexes, and tissues. Timings for embryogenesis show the time post-fertilisation in hours. (**C**) Schematic of life cycle of *Strongyloides stercoralis*. ‘OUT’ denotes aerobically-respiring free-living stages; ‘IN’ indicates stages living inside the host. Note that egg, L1, L2 and L3 can either develop inside or outside the host. (**D**) Graph indicates the percentage of all COQ-2 transcripts that include the RQ-specific exon (‘Percent e-form’) in L3 larvae that either developed outside the host (‘OUT’) or inside the host (‘IN’). Data derive from three individual replicates.

The analysis in *Ascaris* is complicated by the life cycle: while we are comparing *coq-2* splicing in two distinct environments, this is also necessarily a comparison between developing embryos and larvae and adults. It is possible that the changes in splicing we see are not environmentally-induced by the switch from normoxia to anaerobic conditions, but is simply a developmentally programmed switch. To address this, we turned to *Strongyloides stercoralis*. The life cycle of *S. stercoralis* (schematic Figure 6C) is broadly similar to *Ascaris* — L1-L3 stages are free-living, the L3 infective stage infects hosts, and L4 larvae and adults develop and live in the host. However, they have an alternative life cycle where instead of L1-L3 developing outside the host, the eggs can hatch in the host and the entire life cycle takes place inside the host. This allows us to compare the same larval stage in two conditions — here we compare L3 animals that developed inside the host anaerobic environment with L3 animals that developed outside the host. The difference is clear: <20% of *coq-2* transcripts are *coq-2e* in the free-living L3s but >60% is the RQ-synthesizing *coq-2e* form in L3s that developed inside the host (Figure 6D) (Stoltzfus et al., 2012).

In summary, our data show that eukaryotic species that make RQ regulate the choice between making UQ and RQ by alternative splicing of the polyprenyltransferase COQ-2. A switch between two mutually exclusive exons changes the core of the COQ-2 enzyme and switches it from generating UQ precursors to RQ precursors. This alternative splicing event is only seen in species that make RQ and the switch correlates with the change from aerobic to anaerobic metabolism in parasitic helminths.

## Discussion

Organisms are continually challenged by changes in their environments and they must be able to respond for them to survive. Hypoxia is one such challenge and animals have evolved diverse strategies to alter their metabolism to cope with low oxygen levels. For example, humans rapidly switch to anaerobic glycolysis, generating lactate; goldfish on the other hand can adapt to hypoxia by fermenting carbohydrates to generate ethanol (Shoubridge, Hochachka, 1980). In this paper, we focus on how helminths can survive in anaerobic conditions by switching from ubiquinone (UQ)-dependent aerobic metabolism to rhodoquinone (RQ)-dependent anaerobic metabolism. The ability to use RQ is highly restricted amongst animal species — only helminths, molluscs, and annelids are known to make and use RQ. This is a key adaptation since it allows them to rewire their mitochondrial electron transport chain (ETC) to use a variety of terminal electron acceptors in the place of oxygen. They can therefore still use Complex I to pump protons and generate the proton motive force need to power the F0F1ATPase in the absence of oxygen. This allows them to survive without oxygen for long periods — they are thus facultative anaerobes. This ability is critical for parasitic helminths which survive for long periods in the anaerobic environment of the human gut. Since the host does not make RQ or use RQ-dependent metabolism if we could interfere with RQ synthesis, this would be an excellent way to target the parasite and leave the host untouched.

We previously showed that the key decision on whether to make UQ to power aerobic metabolism or RQ to make anaerobic metabolism is dictated by the choice of substrate of the polyprenyltransferase COQ-2 (Roberts Buceta et al., 2019, Del Borrello et al., 2019). COQ-2 must switch from using 4HB to make UQ in aerobic conditions to 3HA to make RQ in anaerobic conditions. Here we reveal the simple mechanism for that switch in substrate specificity in helminths: they use the mutually exclusive alternative splicing of two internal exons to remodel the core of COQ-2. Inclusion of exon 6a results in the COQ-2a enzyme that can make UQ but not RQ; switching 6a for the alternative exon 6e yields COQ-2e which principally makes RQ (Figure 7). All eukaryotes make a homologue of COQ-2a — only the species known to make RQ (helminths, annelids, and molluscs) have genomes encoding the RQ-specific exon 6e and this exon is introduced by alternative splicing in a similar mutually exclusive splicing event in all these phyla. These different lineages thus have the same solution to the problem of substrate switching in COQ-2 — to have two distinct forms of COQ-2 due to alternative splicing — and all do it with the same structural switch in COQ-2.

**Figure 7.**
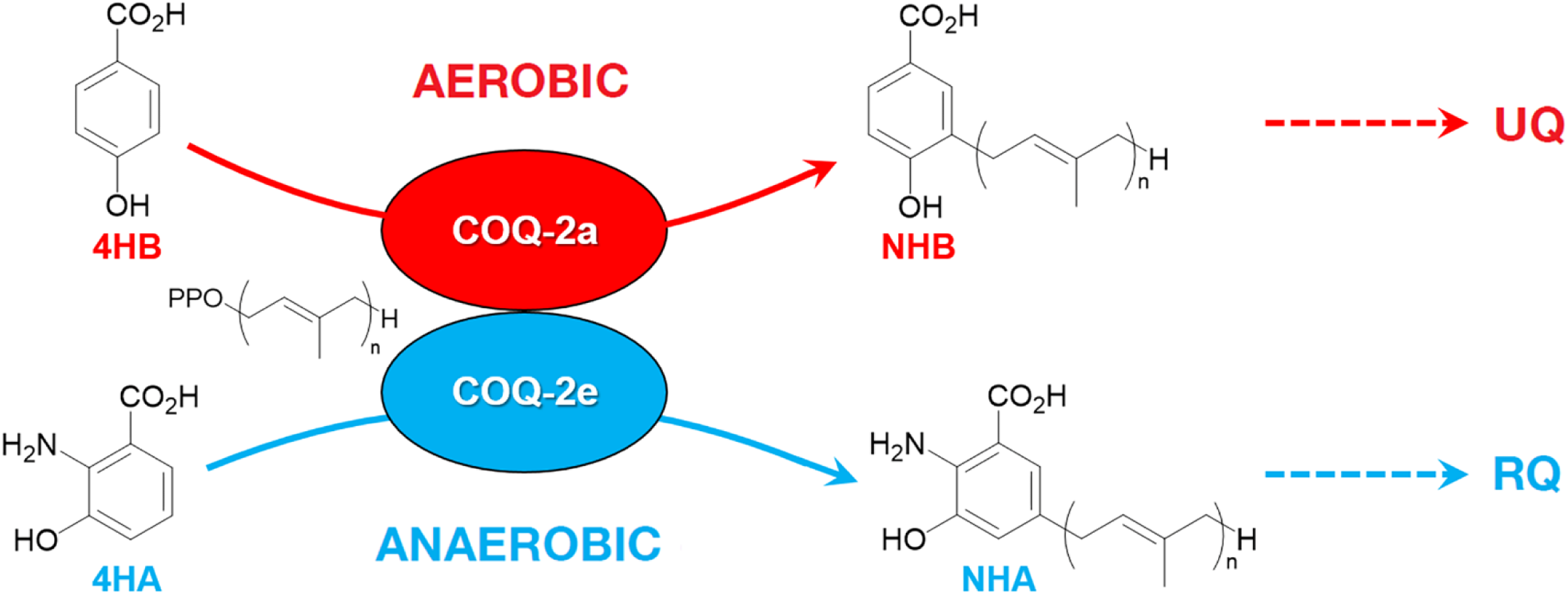
Discrimination between RQ and UQ biosynthesis in *C. elegans*. There are two variants of exon 6 in the *C. elegans coq-2* gene (6a and 6e) which undergo mutually exclusive alternative splicing leading to COQ-2a and COQ-2e isoforms, respectively. Synthesis of UQ originates from 4-hydroxybenzoic acid (4HB) and prenylation is catalyzed by COQ-2a (and marginally by COQ-2e) to form 4-hydroxy-3-nonaprenylbenzoic acid (NHB). In contrast, RQ is most likely synthesized from 3HA, and prenylation is facilitated by COQ-2e to form NHA. Several additional steps are required to convert NHB to UQ and NHA to RQ, respectively.

The alternative splicing of COQ-2 in all lineages that make RQ draws focus to COQ-2 as a potential anthelmintic target. COQ-2e is required for RQ generation and has a distinct sequence to the pan-eukaryotic COQ-2a. The switch between COQ-2a to COQ-2e causes a change in the core of the COQ-2 active site and all species that make RQ have a pair of conserved residues in COQ-2e. We suggest that small molecule inhibitors that selectively target COQ-2e and not the host COQ-2 enzyme could be potent anthelmintics and our groups are initiating these screens at this point. We also note that it is rare to see such a clear and profound change in enzyme specificity due to a single splice event affecting the core of an enzyme catalytic site. There are other examples, most notably a switch in the substrate specificity of the cytochrome P450 CYP4F3 from LTB4 to arachidonic acid (Christmas et al., 1999, Christmas et al., 2001). This switch is tissue specific rather than environmentally induced, however, and there are few other examples to our knowledge. The regulation of COQ-2 specificity by alternative splicing is thus a beautiful and rare example of this type of enzyme regulation by alternative splicing. We note that while alternative splicing of COQ-2 appears to be the key regulated step in determining RQ or UQ biosynthesis, we see no such splicing regulation for other enzymes that are required for both RQ and UQ synthesis downstream of COQ-2 (Roberts Buceta et al., 2019) e.g. there are no known splice variants of COQ-3 and COQ-5, which are quinone methylases downstream of COQ-2. This suggests that while COQ-2 can clearly discriminate between substrates that have/lack a 2-amino group, COQ-3 and COQ-5 would be more promiscuous than COQ-2 and act on both RQ and UQ precursors.

Finally, the mechanism of alternative splicing to regulate the synthesis of UQ or RQ may explain some part of the evolution of RQ synthesis in animals and its phylogenetic distribution. One of the enduring mysteries of RQ synthesis is that it only occurs in very restricted animal phyla. RQ synthesis could either be the ancestral state with widespread loss across most animal species or else RQ synthesis could have arisen independently in helminths, annelids, and molluscs. We believe that the alternative splicing switch from UQ to RQ synthesis favours this independent evolution model. First, we note that there are only two key conserved residues in the RQ-specific exons — this minor sequence change could easily have arisen independently in multiple lineages. Second, there is a clear precedent for a mutually exclusive splicing event that has arisen independently in helminths, annelids, and molluscs in the gene *mrp-1* (Yue et al., 2017). This encodes a transporter that is required for the uptake of vitamin B12 into mitochondria. We note that B12 is critical for aspects of RQ-dependent metabolism. The B12-dependent propionate breakdown pathway (Bulcha et al., 2019) may be required under production of high levels of succinate (under anaerobiosis); succinate can be converted into propionyl-CoA and further used to synthesise branched-chain fatty acids such as 2-methyl butyrate and 2-methyl valerate. This synthesis of branched-chain fatty acids is an RQ-dependent reductive process: RQ carries electrons to ETFDH which is driven in reverse to fatty acid oxidation (Muller et al., 2012). The *mrp-1* gene undergoes alternative splicing in these species and just like in *coq-2, mrp-1* has mutually exclusive exons of exactly the same length. Crucially, *mrp-1* mutually-exclusive alternative splicing has arisen independently in each lineage in which it is seen. We suggest that the reason that RQ synthesis is only seen in helminths, annelids, and molluscs is that these phyla have likewise independently evolved a gene structure of *coq-2* that allows them to switch from a UQ-specific form to a RQ-specific for by mutually-exclusive alternative splicing. Our data thus show that alternative splicing of COQ-2 provides a simple switch from UQ synthesis to RQ synthesis but may also explain how RQ synthesis has arisen independently in multiple distinct animal lineages.

## Materials and methods

### Sequence identification and analysis

To identify COQ-2 sequences from lineages known to synthesize RQ we searched and analyzed from genomes and transcriptomes platyhelminths (*Schistosoma mansoni, Fasciola hepatica* and *Schmidtea mediterranea*), nematodes (*Ascaris suum, Brugia malayi, Haemonchus contortus, Trichuris muris, Strongyloides stercoralis, Strongyloides ratti, Ancylostoma ceylanicum* and *C. elegans*), mollusc (*Crassostrea virginica*, and *Biomphalaria glabrata*) and annelid (*Capitella teleta*). Sequences were retrieved from https://parasite.wormbase.org (WBPS14), *S. mediterranea* database (http://smedgd.neuro.utah.edu), NCBI protein and nucleotide databases and UniProt (https://www.uniprot.org). Human, mice and *Saccharomyces cerevisiae* COQ-2, eukaryotic lineages known to be unable to synthesized RQ, were also identified for comparison. Searches were performed initially with BLASTP (protein databases) using human and *C. elegans* COQ-2 sequences as queries. Additionally, TBLASTN searches were performed using genomic sequences and cDNAs databases. This served to confirm the annotated protein sequences and to identify the non-annotated ones. Identified sequences were confirmed by best reciprocal hits in BLAST. Multiple sequence alignments were made with MUSCLE 3.8 (Chojnacki et al., 2017). Gaps were manually refined after alignment inspection.

### RNA-seq analysis of mutually exclusive exons

To confirm if *coq-2* exons are spliced in a mutually exclusive manner in other helminths, molluscs and annelids, we analyzed existing RNA-seq data for evidence of alternative splicing (listed in Supp. Table 3). Whippet (v0.11) (Sterne-Weiler et al., 2018) was used to analyze RNA-seq data for quantification of AS events. To create a splicing index of exon-exon junctions in Whippet, genome annotations were taken from WormBase Parasite (WBPS14) and Ensembl Metazoa (Release 45). To identify novel exons and splice sites, reads were first aligned to the genome using HISAT2 (Kim, Langmead & Salzberg, 2015). The BAM file generated was then used to supplement existing genome annotations to create a splicing index of known and predicted exon-exon junctions in Whippet (using the --bam --bam-both-novel settings). Where required, TBLASTN data was also used to guide manual re-annotation of the *coq-2* gene. Quantification of AS events was then performed by running whippet quant at default settings. This analysis was repeated for all species listed in Figure 5. A summary of *coq-2* exons with reads that mapped to alternative exon-exon junctions are listed in Supplemental Table 3. We also identified cases where both *coq-2* exons were either included or skipped. However, since these are likely to be non-productive transcripts due to a pre-mature termination codon, we expressed exon usage as the proportion of events where either only the ‘a’ or the ‘e’ form is included.

### Structural analysis

Multiple sequence alignment was performed using Clustal Omega (Madeira et al., 2019). The substrate-bound structure of a UbiA homolog from *A. pernix* (PDB: 4OD5) was displayed on Chimera (Pettersen et al., 2004) and the *C. elegans* sequence was threaded by homology using Modeller (Sali, Blundell, 1993, Webb, Sali, 2016).

### *Caenorhabditis elegans* strains and culture conditions

The *C. elegans* wild-type Bristol strain (N2) was obtained from the Caenorhabditis Genetics Center (CGC, University of Minnesota, USA), which is supported by the National Institutes of Health-Office of Research Infrastructure Programs. The *C. elegans* mutant strains in *coq-2* exon 6A (PHX1715, *coq-2(syb1715)*) and *coq-*2 exon 6E (PHX1721, *coq-2(syb1721)*) were generated by Suny Biotech Co., Ltd (Fuzhou City, China) using CRISPR/Cas9 system. The precise deletion of both mutant strains (134 bp) was verified by DNA sequencing the flanking region of exons 6a and 6e. The wild-type sequence, the deleted sequence in each strain, the sgRNAs, and primers used are listed in Supplemental Table 1.

The general methods used for culturing and maintenance of *C. elegans* are described in (Brenner, 1974). All chemical reagents were purchased from Sigma-Aldrich (St. Louis, MO, USA). The *E. coli* OP50 strain, used as *C. elegans* food, was also received from CGC.

### Lipid extraction and LC-MS quantitation

Lipid extractions of *C. elegans* N2 and mutant strains were performed on ∼100 mg worm pellets after adding 1000 pmol UQ3 internal standard (11). LC-MS samples were prepared from lipid extracts and diluted 1:100 (Bernert, et *al.*, 2019). Standards were extracted using the same method as for worm samples at the following concentrations: UQ_3_ (10 pmol/10 µL injection), RQ_9_ (0.75, 1.5, 3.0, 4.5, or 6.0 pmol/10 µL injection), and UQ_9_ (3.75, 7.5, 15.0, 22.5, or 30.0 pmol/10 µL injection). The Q_3_ standard was synthesized at Gonzaga University (Campbell et al., 2019), the RQ_9_ standard was isolated by preparative chromatography from *A. suum* lipid extracts (Roberts Buceta et al., 2019) and the UQ_9_ standard was purchased (Sigma-Aldrich, St. Louis, MO). The general LC-MS conditions and parameters were previously reported (Bernert et al., 2019, Campbell et al., 2019). Samples were analyzed in quadruplicate and the pmol quinone was determined from the standard curve and corrected for recovery of internal standard. Samples were normalized by mg pellet mass.

### Image-based KCN recovery assay

The KCN recovery assay was performed as previously described (Spensley et al., 2018, Del Borrello et al., 2019). Briefly, L1 worms were isolated by filtration through an 11 µm nylon mesh filter (Millipore: S5EJ008M04). Approximately 100 L1 worms in M9 were dispensed to each well of a 96 well plate and an equal volume of potassium cyanide (KCN) (Sigma-Aldrich St. Louis, MO) solution was then added to a final concentration of 200 µM KCN. Upon KCN addition, plates were immediately sealed and incubated at room temperature for 15 h on a rocking platform. After 15 h, the KCN was diluted 6-fold by addition of M9 buffer. Plates were immediately imaged on a Nikon Ti Eclipse microscope every 10 min for 3 h. Fractional mobility scores (FMS) were then calculated using a custom image analysis pipeline (Spensley et al., 2018). For each strain, FMS scores for the KCN-treated wells were normalized to the M9-only control wells at the first timepoint. Three technical replicates were carried out in each experiment and the final FMS scores taken from the mean of four biological replicates.

## Acknowledgements

*C. elegans* strain N2, and *E. coli* OP50 were provided by the Caenorhabditis Genetics Center, which is funded by NIH Office of Research Infrastructure Program: P40 OD010440. We thank Exequiel Barrera, John Calarco and Amy Caudy for helpful discussions.

## Supplementary Documentation

**SUPPLEMENTAL TABLE 1.**
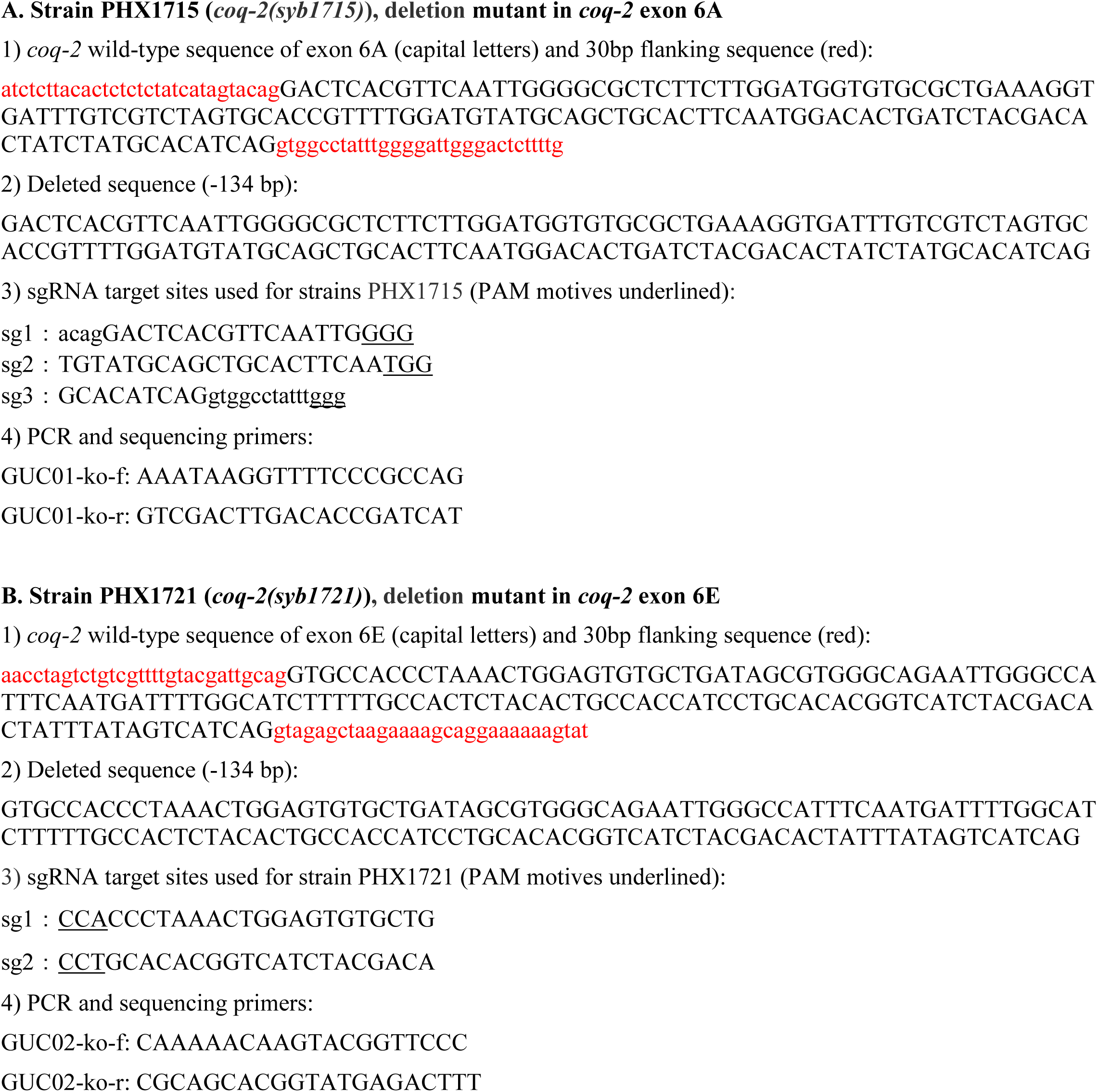
*C. elegans* strains.

**SUPPLEMENTAL TABLE 2.** Statistical analysis of RQ_9_ and Q_9_ levels in *coq-2* mutant strains

| Strain | Avg pmol<br>RQ<sub>9</sub>/mg pellet | p value<br>(N = 4) | Avg pmol<br>Q<sub>9</sub>/mg pellet | p value<br>(N = 4) |
| --- | --- | --- | --- | --- |
| N2 | 3.30 ± 0.74 |  | 16.55 ± 4.76 |  |
| *coq-2*Δ6A | 5.43 ± 1.35 | 0.016 | 2.81 ± 0.27 | < 0.001 |
| *coq-2*Δ6E | 0.40 ± 0.08 | < 0.001 | 20.23 ± 3.54 | 0.130 |

**SUPPLEMENTAL TABLE 3.**
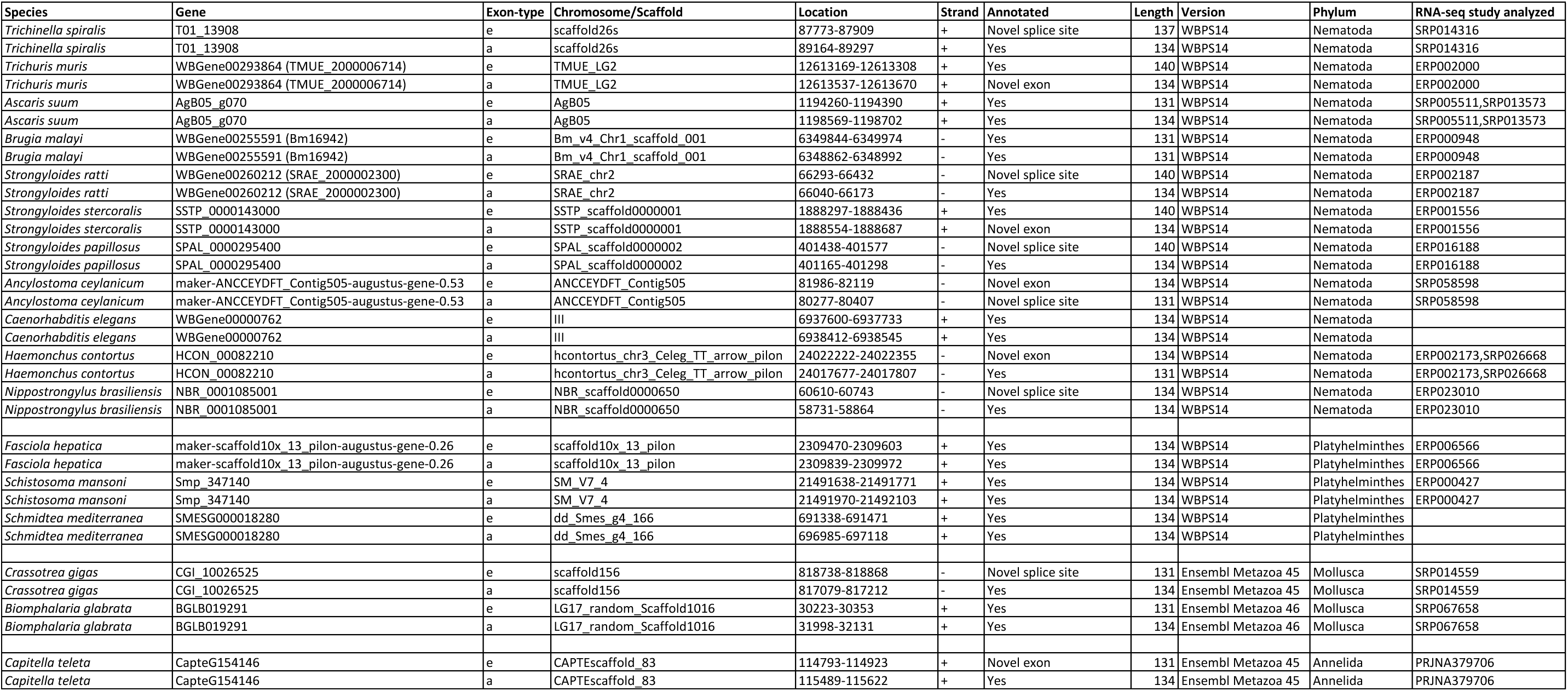
Genomic coordinates of known and predicted mutually exclusive *coq-2* a/e exons

